# MANF Clears Mutant Uromodulin in Human Kidney Organoids of Autosomal Dominant Tubulointerstitial Kidney Disease

**DOI:** 10.64898/2026.03.02.708095

**Authors:** Chenjian Gu, Yili Fang, Yixuan Wang, Eric Tycksen, Gayathri Kondepati, Chuang Li, Kendrah Kidd, Jun Liu, Fumihiko Urano, Maria Lindahl, Anthony J. Bleyer, Srikanth Singamaneni, Zhao Sun, Ying Maggie Chen

## Abstract

Autosomal dominant tubulointerstitial kidney disease due to uromodulin mutations (ADTKD-UMOD) is one of the leading hereditary kidney diseases. Currently there is no targeted treatment. To illuminate human relevance of mesencephalic astrocyte-derived neurotrophic factor (MANF)-based therapy, we have established patient induced pluripotent stem cell (iPSC)-derived kidney organoid model carrying UMOD p.H177-R185del, the leading mutation causing ADTKD. We have discovered that MANF can directly bind and repress ER calcium release channel IP3R1, thus enhancing AMPK-induced autophagy in a TRIB3-dependent manner. The therapeutic implication of this finding may well be extended to other protein misfolding diseases.

---

Autosomal dominant tubulointerstitial kidney disease due to *UMOD* mutations (ADTKD-*UMOD*) is a leading monogenic cause of chronic kidney disease (CKD), and *UMOD* p.H177-R185del is the most prevalent pathogenic variant in the US, accounting for 21% of ADTKD-*UMOD*. Currently, therapy is unavailable. UMOD is synthesized and secreted by thick ascending limb (TAL) of Henle’s loop. We generated the first ADTKD mouse model carrying the human H177-R185del-equivalent deletion mutation and demonstrated that ER stress triggered by the mutant UMOD suppresses p-AMPK-FOXO3-mediated autophagy in TALs. Moreover, overexpression of mesencephalic astrocyte-derived neurotrophic factor (MANF), an ER chaperone, ameliorates CKD by enhancing p-AMPK in mice (1).

To further bolster up translational relevance and delineate how MANF activates p-AMPK, induced pluripotent stem cells (iPSCs) from a heterozygous H177-R185del (*UMOD* ^DEL/+^) patient and an unaffected family member (Figure 1A) were successfully differentiated to kidney organoids expressing UMOD. The under-glycosylated mutant UMOD localized to the ER with diffuse intra-tubular expression, whereas WT UMOD displayed apical TAL enrichment (Figure 1, B and C). Furthermore, for the first time, we isolated TALs from organoids. *UMOD* ^DEL/+^ TALs recapitulated hallmark features of our ADTKD mice, including ER stress activation, manifested by BiP and p50ATF6 induction, as well as autophagy impairment, characterized by p-AMPK/FOXO3 inhibition and p62 and LC3B-II accumulation (Figure 1D). In the mutant TALs, RNAseq showed upregulation of TAL injury markers, as identified in the Human Kidney Atlas (HKAv2) (2), and KEGG analysis revealed inflammation as the most upregulated pathway (Supplemental Figure 1, A and B).

**Figure 1.**
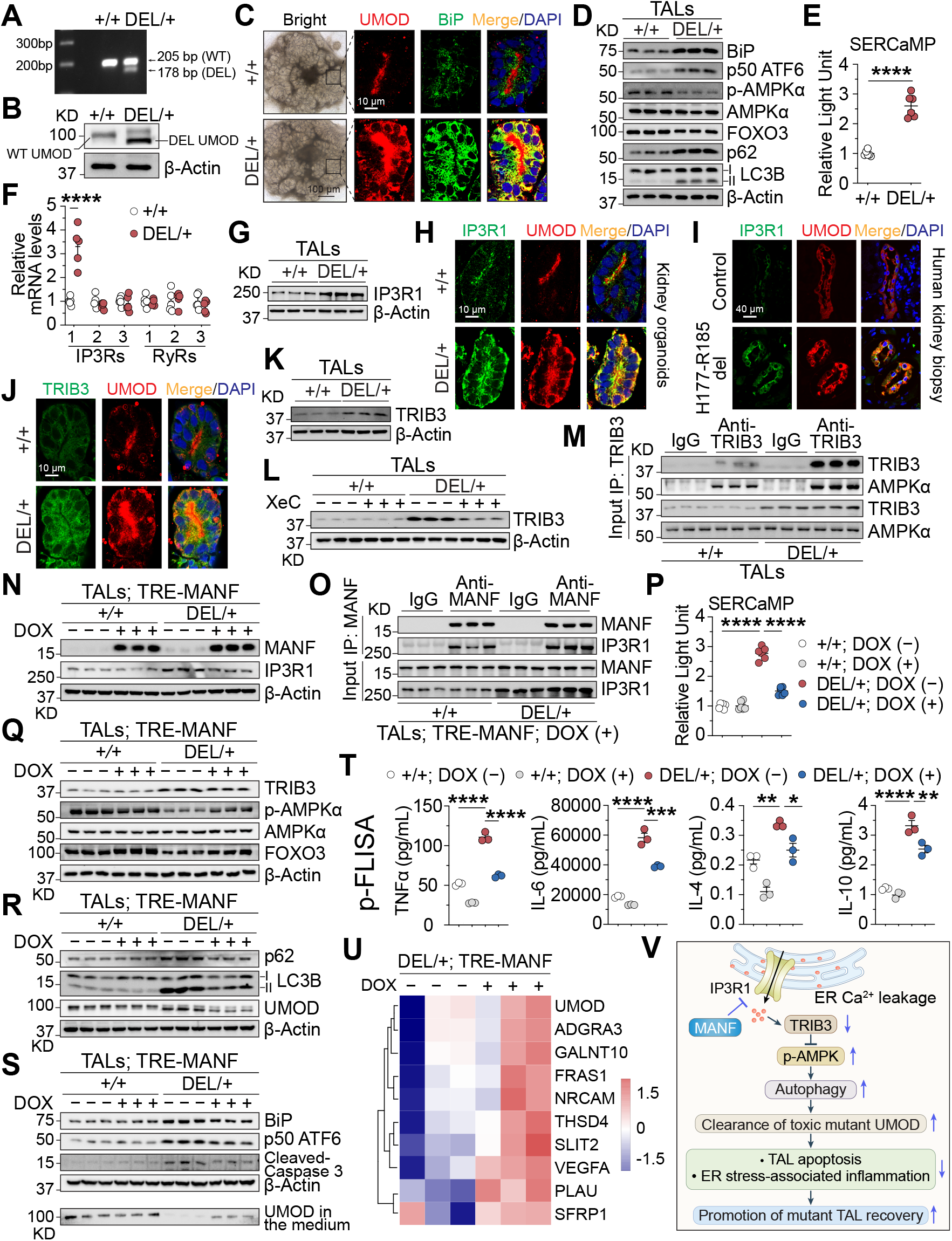
MANF promotes UMOD proteostasis by regulating IP3R1-mediated ER calcium efflux in ADTKD kidney organoids. (**A**) Genotyping of hiPSCs from a WT and heterozygous patient with *UMOD* p.H177-R185del. (**B**) WB of UMOD in organoids. (**C**) Bright-field or double-IF of UMOD, BiP and DAPI on WT or mutant organoids. (**D, G, K**) IB of indicated proteins in organoid-isolated WT or DEL/+ TALs. (**E**) Luciferase activity in the organoid medium. (**F**) Transcript expression of ER calcium efflux channels in organoid-derived TALs. (**H–I**) IF of IP3R1 and UMOD on organoids or human kidney sections. (**J**) Dual-IF staining of TRIB3 and UMOD on organoids. (**L**) TAL IB of TRIB3 without or with XeC treatment. (**M**) Co-IP demonstrating the association of TRIB3 and AMPKα in organoid-derived TALs. (**N–U**) WT or DEL/+ organoids were treated without or with DOX from days 22**–**30. (**N, Q–S**) IB of the indicated proteins in organoid-derived TALs. (**O**) Co-IP showing the interaction of MANF and IP3R1 in TALs with DOX. (**P**) Luciferase activity in the organoid medium. (**T**) p-FLISA of cytokines in the organoid medium. (**U**) Heatmap of TAL healthy state markers. Benjamini-Hochberg adjusted *P*<0.05. (**V**) Summary illustrating the proposed protective mechanisms of MANF in ADTKD-*UMOD* organoids. \**P*<0.05; \*\**P*<0.01; \*\*\**P*<0.001; \*\*\*\**P*<0.0001.

Given that ER stress and calcium dynamics are intricately linked, novel secreted ER-calcium-monitoring protein (SERCaMP) (3)-expressing WT or DEL/+ hiPSCs were utilized, and differentiated DEL/+ organoids exhibited marked ER calcium leakage (Figure 1E). Detailed examination of key ER calcium release channels revealed selective increase of inositol 1,4,5-trisphosphate receptor 1 (IP3R1) at both transcriptional and translational levels, but not upregulation of IP3R2, IP3R3 and ryanodine receptors (RyRs) in the mutant TALs (Figure 1, F– H), aligning with the human kidney biopsy finding (Figure 1I). We next sought to investigate whether tribbles homolog 3 (TRIB3), an ER stress-inducible pseudokinase, is implicated following ER calcium leak. Dual-immunofluorescence (IF) staining and IB of organoid-derived TALs showed that the pronounced induction of TRIB3 in DEL/+ TALs (Figure 1, J and K) was attenuated by the IP3R blocker Xestospongin C (XeC) (Figure 1L), indicating TRIB3 as a downstream effector of ER calcium deletion.

Since AMPK serves as a key linker between stress responses and autophagy promotion, we further determined whether TRIB3 could modulate AMPK activity. Co-IP revealed that endogenous TRIB3 directly binds to the catalytic subunit AMPKα in the organoid-isolated TALs (Figure 1M). To further map the TRIB3 interaction region, the deletion mutants of FLAG-tagged AMPKα1 or AMPKα2 were constructed. Pull-down assays demonstrated a direct interaction between TRIB3 and the activation loop (AL) domain of both AMPKα1 and AMPKα2 (Supplemental Figure 2, A–C), which may block T172/T183 phosphorylation within the AL. In stable HEK293 cells expressing WT or *UMOD* H177-R185del, TRIB3 was induced and p-AMPKα (T172/T183) was reduced in DEL vs. WT cells as well (Supplemental Figure 2E). Overexpression and knockdown of TRIB3 further validated that TRIB3 negatively regulated AMPKα phosphorylation (Supplemental Figure 2, D and E). Collectively, these data demonstrate that in ADTKD organoids, IP3R1-mediated ER calcium deletion, triggered by misfolded UMOD-induced ER stress, upregulates TRIB3, which can directly bind to and suppress AMPK activation, thereby impairing autophagy.

To determine whether MANF exerts a therapeutic role in ADTKD organoids, tetracycline-inducible MANF-overexpressing (TRE-MANF) iPSCs were generated (Supplemental Figure 3A) and differentiated into kidney organoids. Upon doxycycline induction from Days 22–30, MANF overexpression in the DEL/+ organoids, including TALs (Figure 1N and Supplemental Figure 3B) significantly mitigated IP3R1 expression through direct interaction with IP3R1 (Figure 1, N and O), thus attenuating ER calcium depletion (Figure 1P). The subsequent TRIB3 repression and restoration of p-AMPK/FOXO3 expression by MANF upregulation (Figure 1Q) promoted autophagic flux (Supplemental Figure 3C) and clearance of mutant UMOD (Figure 1R), leading to ER stress and apoptosis amelioration, as well as enhanced WT UMOD secretion to the organoid medium (Figure 1S). Additionally, our newly invented, ultrasensitive plasmon-enhanced fluorescence-linked immunosorbent multiplex assay (p-FLISA) (4) (Supplemental Figure 3D) showed that MANF overexpression inhibited pro-inflammatory cytokine secretion to the medium (Figure 1T) and their concordant synthesis (Supplemental Figure 3E) in the mutant organoids. Consequently, enhanced MANF expression in the mutant organoids led to mutant TAL recovery, as demonstrated by increased expression of healthy TAL genes (HKAv2) (Figure 1U).

Importantly, MANF-overexpressing lentivirus treatment of DEL/+ organoids, from Days 22– 30, blocked IP3R1-mediated ER calcium depletion and subsequently relieved TRIB3-induced p-AMPK suppression and defective autophagic degradation of mutant UMOD (Supplemental Figure 4, A–E). Correspondingly, the lentivirus treatment reproduced the protective effects of lentiviral vector-mediated MANF upregulation in DEL/+ hiPSCs (Supplemental Figure 4, F–H).

In summary, we have discovered a novel therapeutic mechanism, whereby MANF restores compromised autophagy by blunting ER-calcium-efflux in ADTKD organoids (Figure 1V). By directly targeting IP3R1, MANF may emerge as a potent therapy capable of correcting disrupted proteostasis for ADTKD, and potentially for a spectrum of protein misfolding diseases. Additionally, our ADTKD organoids, as a human-relevant model, provide a powerful high-throughput drug screening platform for identifying more therapeutics, including human RNA-based therapy.

