## Supplemental Material for "MANF Clears Mutant Uromodulin in Human Kidney Organoids of Autosomal Dominant Tubulointerstitial Kidney Disease"

### Methods

#### Cell culture, transfection, and generation of stable cell lines

Human embryonic kidney cells (HEK 293T and HEK 293; ATCC) were maintained in Dulbecco's modified Eagle's medium (DMEM; Gibco), supplemented with 10% fetal bovine serum (FBS; Gibco) and antibiotics (50 U/mL penicillin and 50 µg/mL streptomycin; Gibco). Cells were cultured at 37 °C in humidified incubator with 5% $CO_2$. Transient transfection of plasmids into HEK 293T cells or HEK 293 cells was performed using Lipofectamine™ 3000 Transfection Reagent (Invitrogen) following the manufacturer's instructions. Stable HEK 293 cell lines expressing UMOD or UMOD-H177-R185 del were previously described (1).

Human-induced pluripotent stem cells (hiPSCs) were generated through reprogramming dermal fibroblasts from an ADTKD patient harboring the *UMOD* p.H177-R185 deletion mutation, as well as from an unaffected family member. This reprogramming was carried out at Washington University Genome Engineering and iPSC Center (WU GEiC). hiPSCs were maintained under feeder-free conditions on Matrix-coated plates (Corning) in mTeSR Plus medium (STEMCELL Technologies). Cells were passaged every 3-4 days using ReLeSR (STEMCELL Technologies) and routinely tested for mycoplasma contamination. hiPSCs generated in this study were examined for their karyotypes by WU GEiC to ensure genomic integrity.

For tetracycline (Tet)-inducible MANF, cytomegalovirus (CMV) promoter driven MANF, or phosphoglycerate kinase promoter (PGK) driven SERCaMP, lentiviral particles were produced by transient co-transfection of HEK 293T cells with respective transfer plasmid (pLVX-TRE (tetracycline-responsive element) 3G-MANF-PGK-rtTA (reverse tetracycline-controlled transactivator)-P2A-Blast, pLVX-CMV-MANF or pLenti6.3-PGK-SERCaMP), packaging plasmid psPAX2 (Addgene), and envelope plasmid pMD2.G (Addgene). Culture medium was replaced 8 hours post-transfection, and viral supernatants were collected at 48 and 72 hours, then filtered through a 0.45 μm syringe filter and applied to target hiPSCs. Stably transduced hiPSCs were selected with 0.5 μg/mL Blasticidin (Gold Biotechnology) or 2 μg/mL Puromycin (Sigma-Aldrich) and maintained under standard hiPSC culture conditions.

**Plasmid construction**

For Tet-inducible MANF expression, the lentiviral vector pLVX-TRE3G-MANF-PGK-rtTA-P2A-Blast was constructed using Gibson Assembly cloning method (NEB). In this inducible system, human *MANF* sequence (NM_006010.6) was cloned downstream of the TRE3G promoter, which confers tight and doxycycline-responsive transcriptional control, enabling doxycycline-inducible transcription. To achieve simultaneous expression of the rtTA and the Blasticidin resistance gene from a single transcript, a P2A self-cleaving peptide sequence was inserted between the rtTA and Blasticidin resistance gene coding sequences. This design facilitates efficient co-expression by ribosomal skipping, ensuring balanced production of both proteins without requiring an internal ribosome entry site (IRES). The rtTA transactivator and Blasticidin resistance cassette were placed under the control of the constitutive PGK promoter, providing stable and consistent expression for selection purposes. All DNA fragments including the TRE3G promoter, *MANF* gene, PGK promoter, rtTA, P2A, and Blasticidin resistance gene, were PCR amplified with primers containing overlapping sequences compatible with Gibson Assembly. The assembled plasmid was verified by Sanger sequencing (GENEWIZ from Azenta Life Sciences) to confirm correct orientation and sequence integrity.

The pLVX-CMV-MANF construct was generated by cloning the human MANF coding sequence into the pLVX vector under the control of the CMV promoter. The pLenti6.3-PGK-SERCaMP plasmid was generated by replacing the CMV promoter in the original pLenti6.3-SERCaMP construct with the PGK promoter. The pLenti6.3-SERCaMP plasmid was kindly provided by Dr. Harvey (2).

The construct pcDNA3.1-HA-TRIB3, encoding N-terminally HA-tagged human TRIB3, was generated using pcDNA3.1-HA as the backbone vector by inserting the human TRIB3 coding sequence (NM_001301188.1) in-frame with the HA tag. The expression plasmids pcDNA3.1-FLAG-AMPKα1 and pcDNA3.1-FLAG-AMPKα2, encoding N-terminally FLAG-tagged human AMPKα1 and AMPKα2, were generated by cloning *PRKAA1* (NM_001355028.2) and *PRKAA2* (NM_006252.4), respectively, into the pcDNA3.1-FLAG vector, in-frame with the FLAG tag. Full-length human TRIB3, PRKAA1, and PRKAA2 cDNA were amplified by PCR from HEK293T total cDNA pools. Truncation mutants of AMPKα1 and AMPKα2 were generated via site-directed mutagenesis using the KOD-Plus-Mutagenesis Kit (TOYOBO). All constructs were confirmed by Sanger sequencing.

Lentiviral vector pLKO.1 (Addgene) was used for generating shRNAs targeting TRIB3. Target sequences were selected as follows: shTRIB3#1, 5′-GATCTCAAGCTGTGTCGCTTT-3′; shTRIB3#2, 5′-GCCGTGCTCTTCCGCCAGATG-3′. A non-targeting scrambled shRNA 5′-CCTAAGGTTAAGTCGCCCTCG-3′ was used as a control.

**Generation of hiPSC-derived 3D kidney organoids**

Kidney organoids were generated following a previously published protocol (3). Briefly, 375,000 human iPSCs were seeded into T25 flasks in mTeSR Plus medium (STEMCELL Technologies), supplemented with 10 μM Rho-associated kinase (ROCK) inhibitor Y-27632 (Tocris Bioscience). After 12 hours, the medium was replaced with APEL2 medium (STEMCELL Technologies) containing 8 μM CHIR99021 (Tocris Bioscience). Cells were maintained in this medium for 4 days, followed by a 3-day incubation in APEL2 medium supplemented with 200 ng/mL FGF9 (R&D Systems) and 1 μg/mL heparin (Sigma-Aldrich). On day 7, cells were dissociated into single cells using Accutase cell dissociation reagent (Gibco). A total of 500,000 cells were pelleted by centrifugation at 400 × g for 5 minutes and transferred onto Transwell with 0.4 μm pore polyester membrane inserts (Corning). Pellets were first treated with 5 μM CHIR99021 in APEL2 medium for 1 hour and then cultured for 5 days in APEL2 medium containing 200 ng/mL FGF9 and 1 μg/mL heparin. For the next 18 days, organoids were maintained in APEL2 medium supplemented with 1 μg/mL heparin, which was refreshed every other day.

Large numbers of kidney micro-organoids were generated via a suspension culture approach with minimal modifications (4). In brief, 375,000 iPSCs were seeded in T25 flasks with mTeSR Plus medium containing 10 μM Y-27632. After 12 hours, cells were treated with 7 μM CHIR99021 in APEL2 medium for 4 days, followed by APEL2 medium supplemented with 200 ng/mL FGF9, 1 μM CHIR99021, and 1 μg/mL heparin for an additional 3 days. On day 7, cells were enzymatically dissociated into single cells using Accutase and seeded into round-bottom, ultra-low attachment 96-well plates (Corning) at 100,000 cells per well in APEL medium supplemented with 200 ng/mL FGF9, 1 μM CHIR99021, 0.1% polyvinyl alcohol (PVA; Sigma-Aldrich), 0.1% methylcellulose (MC; Sigma-Aldrich), 1 μg/mL heparin, and 10 μM Y-27632. After 24 hours, organoids spontaneously formed within the wells. The medium was then replaced with APEL medium containing 200 ng/mL FGF9, 1 μM CHIR99021, 0.1% PVA, 0.1% MC, and 1 μg/mL heparin and maintained for 4 additional days. Subsequently, organoids were cultured in

APEL2 medium containing 1 μg/mL heparin, with medium changed every other day for up to 18 days.

**MANF overexpression in kidney organoids**

For inducible lentiviral vector-mediated MANF overexpression, TRE-MANF-expressing stable hiPSC line-differentiated organoids were cultured in APEL medium supplemented with 1 μg/mL heparin, and 0.5 μg/mL doxycycline (Sigma-Aldrich) administered from day 22 to day 30. For lentivirus-mediated MANF overexpression, kidney organoids were infected overnight on day 22 with lentivirus encoding human MANF, then cultured in APEL medium supplemented with 1 μg/mL heparin until day 30. All organoids were harvested on day 30 for downstream analysis.

**Immunofluorescence staining of WT and ADTKD kidney organoids**

Organoids were immersion-fixed in 4% paraformaldehyde (PFA) for 20 minutes at room temperature, followed by cryoprotection in 30% (w/v) sucrose in PBS at 4 °C overnight to ensure complete dehydration. The cryoprotected organoids were then embedded in optimal cutting temperature (OCT) compound and snap-frozen using dry ice or an isopentane/liquid nitrogen slurry. For immunofluorescence analysis, embedded organoids were cryosectioned at a thickness of 8 μm using a cryostat and mounted onto Superfrost Plus microscope slides (Thermo Fisher Scientific). Sections were air-dried and subsequently washed three times with PBS to remove residual OCT. Antigen retrieval was performed by heating the sections in citrate buffer (10 mM, pH 6.0) at 95 °C for 20 min, followed by cooling to room temperature. Nonspecific binding sites were blocked by incubating the sections in PBS containing 1% bovine serum albumin (BSA) for 1 hour at room temperature. Primary antibodies were diluted in PBS containing 0.1% Triton X-

100 and 1% BSA and applied to the sections for overnight incubation at 4 °C in a humidified chamber. The following day, sections were washed and incubated with appropriate fluorophore-conjugated secondary antibodies diluted in the same buffer for 1 hour at room temperature, protected from light. Nuclear counterstaining was performed using 4',6-diamidino-2-phenylindole (DAPI) for 10 minutes. Slides were mounted using antifade mounting medium (Invitrogen) and imaged using a Leica TCS SP8 laser-scanning confocal microscope (Leica Microsystems). Image acquisition settings were kept the same across different experimental groups.

**Purification of TAL epithelial cells from kidney organoids**

UMOD-expressing epithelial cells were isolated from differentiated kidney organoids as previously described (1) with minor modifications. Briefly, kidney organoids were enzymatically dissociated into single-cell suspensions using Accutase cell dissociation reagent (Gibco). Organoids were incubated in Accutase at 37 °C with gentle shaking for 5 minutes. Enzymatic digestion was terminated by adding an equal volume of DMEM/F12 (Gibco) supplemented with 10% fetal bovine serum (Gibco). Following digestion, cell suspensions passed through a 40 μm cell strainer (Alkali Scientific) to remove residual clumps, yielding a single-cell suspension suitable for magnetic bead-based cell sorting. UMOD-positive cells were isolated using a magnetic bead-based immunocapture approach. Biotinylated polyclonal sheep anti-human UMOD antibody (R&D Systems) was incubated with Dynabeads Biotin Binder magnetic beads (Invitrogen). For each preparation, 100 μL of Dynabeads were conjugated with 2 μg of anti-UMOD antibody by incubation at 4 °C for 2 hours with gentle rotation. After washing to remove unbound antibody, the conjugated beads were incubated with the single-cell suspension at 4 °C for 1 hour to allow binding to UMOD-expressing cells. Bead-bound UMOD-positive cells were isolated using a magnetic separation stand. The bead-cell complexes were retained against the magnet, and unbound (UMOD-negative) cells were removed through multiple PBS washes. The enriched UMOD-positive fraction was then collected and processed for downstream RNA and protein extraction using standard protocols.

To inhibit $IP_3R1$-mediated $Ca^{2+}$ leakage, isolated TALs were incubated with 400 nM Xestospongin C (XeC; Sigma-Aldrich), a selective $IP_3R$ inhibitor, for one hour prior to downstream analyses.

**mRNA quantification by quantitative real-time PCR**

Total RNA was extracted from isolated TAL cells using the RNeasy UCP Micro Kit (QIAGEN), following the manufacturer's protocol. Genomic DNA contamination was eliminated by on-column DNase I treatment included in the kit. RNA concentration and purity were assessed using a NanoDrop spectrophotometer. For cDNA synthesis, RNA was reverse transcribed using PrimeScript RT Reagent Kit (Takara Bio) according to the manufacturer's instructions. Real-time PCR was performed using Fast SYBR Green Master Mix (Applied Biosystems). Amplification was carried out on a QuantStudio 6 Flex Real-Time PCR System (Applied Biosystems) under the following thermal cycling conditions: initial denaturation at 95 °C for 20 seconds, followed by 40 cycles of denaturation at 95 °C for 1 second and annealing/extension at 60 °C for 20 seconds. Melt curve analysis was performed to verify specificity of amplification. Relative gene expression was calculated using the ΔΔCt method, with *GAPDH* serving as internal normalization controls. Data were analyzed and presented as fold changes relative to control groups.

**RNA sequencing and analysis**

RNA integrity was determined using Agilent Bioanalyzer or 4200 Tapestation. Library preparation was performed with 10ng of total RNA with a Bioanalyzer RIN score greater than 8.0. ds-cDNA was prepared using the SMARTer Ultra Low RNA kit for Illumina Sequencing (Takara-Clontech) per manufacturer's protocol. cDNA was fragmented using a Covaris E220 sonicator using peak incident power 18, duty factor 20%, cycles per burst 50 for 120 seconds. cDNA was blunt ended, had an A base added to the 3' ends, and then had Illumina sequencing adapters ligated to the ends. Ligated fragments were then amplified for 12-15 cycles using primers incorporating unique dual index tags. Fragments were sequenced on an Illumina NovaSeq X Plus using paired end reads extending 150 bases. Basecalls and demultiplexing were performed with Illumina's DRAGEN BCLconvert version 4.3.13. RNA-seq reads were then aligned to the Ensembl GRCh38.113 primary assembly with STAR version 2.7.11b (5). Gene counts were derived from the number of uniquely aligned unambiguous reads by Subread:featureCount version 2.0.8 (6). Isoform expression of known Ensembl transcripts were quantified with Salmon version 1.10.0 (7). Sequencing performance was assessed for the total number of aligned reads, total number of uniquely aligned reads, and features detected. The ribosomal fraction, known junction saturation, and read distribution over known gene models were quantified with RSeQC version 5.0.4 (8).

All gene counts were then imported into the R/Bioconductor v4.5.1 package EdgeR (9) and TMM normalization size factors were calculated to adjust for samples for differences in library size. Ribosomal genes and genes not expressed in at least 3 samples greater than 1 count-per-million were excluded from further analysis. The TMM size factors and the matrix of counts were then imported into the R/Bioconductor package Limma (10). Weighted likelihoods based on the observed mean-variance relationship of every gene and sample were then calculated for all samples with the voomWithQualityWeights (11) function and were fitted using a Limma generalized linear model with additional unknown latent effects as determined by surrogate variable analysis (SVA) (12). The performance of all genes was assessed with plots of the residual standard deviation of every gene to their average log-count with a robustly fitted trend line of the residuals. Differential expression analysis was then performed to analyze for differences between conditions, and the results were filtered for only those genes with Benjamini-Hochberg false-discovery rate adjusted p-values less than or equal to 0.05.

For each contrast extracted with Limma, global perturbations in known Gene Ontology (GO) terms, MSigDb, and KEGG pathways were detected using the R/Bioconductor package GAGE (13) to test for changes in expression of the reported log 2 fold-changes reported by Limma in each term versus the background log 2 fold-changes of all genes found outside the respective term. The R/Bioconductor package heatmap3 (14) was used to display heatmaps across groups of samples for each GO or MSigDb term with a Benjamini-Hochberg false-discovery rate adjusted p-value less than or equal to 0.05. Perturbed KEGG pathways where the observed log 2 fold-changes of genes within the term were significantly perturbed in a single-direction versus background with p-values less than or equal to 0.05 were rendered as annotated KEGG graphs with the R/Bioconductor package Pathview (15).

**Immunoprecipitation**

To immunoprecipitate MANF, TRIB3 or HA-TRIB3 proteins, 2 μg of corresponding antibodies or control normal rabbit IgG (Invitrogen) were incubated with 20 μL of Protein A/G PLUS-Agarose beads (Santa Cruz Biotechnology) at room temperature for 1 hour, with gentle rotation to allow antibody-bead conjugation. After binding, cells were lysed with IP lysis buffer (25 mM Tris-HCl, pH 7.4; 150 mM NaCl; 1% NP-40; 1 mM EDTA; 5% glycerol) containing protease and phosphatase inhibitor cocktails (Sigma-Aldrich), and the antibody-bead complexes were incubated with cell lysates at 4 °C overnight under constant rotation. Following incubation, the beads were washed three times with ice-cold IP lysis buffer to remove non-specifically bound proteins. Immunoprecipitated proteins were eluted by boiling in SDS loading buffer and subjected to downstream analysis by SDS-polyacrylamide gel electrophoresis (SDS-PAGE) and immunoblotting.

**Western blot analysis**

UMOD-positive cells were lysed in RIPA buffer (Cell Signaling Technology) supplemented with protease and phosphatase inhibitor cocktails (Sigma-Aldrich). Lysates were incubated on ice for 10 minutes with periodic vortexing and subsequently centrifuged at 14,000 × g for 15 minutes at 4 °C. The supernatants were collected, and total protein concentrations were determined using the DC Protein Assay Kit (Bio-Rad Laboratories) according to the manufacturer's instructions, with BSA as the standard. Equal amounts of protein were denatured by boiling in SDS loading buffer containing 1% β-mercaptoethanol (Sigma-Aldrich) for 5 minutes and resolved by SDS-PAGE. Proteins were subsequently transferred to polyvinylidene difluoride (PVDF) membranes (Thermo Fisher Scientific). Membranes were blocked in Tris-buffered saline containing 0.1% Tween-20 (TBST) and 5% non-fat milk for 1 hour at room temperature, and then incubated overnight with primary antibodies at 4 °C. After washing three times in TBST, the membranes were incubated for 1 hour at room temperature with the appropriate horseradish peroxidase (HRP)-conjugated secondary antibodies (Cell Signaling Technology). Immunoreactive bands were detected using Clarity Max ECL substrate (Bio-Rad Laboratories) and visualized with the ChemiDoc™ MP Imaging System (Bio-Rad Laboratories). To confirm equal protein loading, the same blot was stripped with stripping buffer (25 mM glycine, 1% SDS, pH 2.0) and then incubated with an HRP-conjugated anti-β-actin antibody (Sigma-Aldrich).

**Human IL-6 ELISA**

IL-6 concentrations in organoid medium were quantified using Human IL-6 DuoSet ELISA kit (R&D Systems). Specifically, 96-well plates were first coated with capture antibodies through overnight incubation at room temperature, followed by blocking with 200 µl of 3% BSA. After washing with PBST (0.05% Tween-20 in PBS), 100 µl of serial diluted IL-6 standard, as well as organoid medium samples (100-fold dilution using 1% BSA in PBS) were added in duplicates and incubated at room temperature for 2 hours. The plate was washed subsequently and incubated with biotinylated detection antibodies (50 ng /ml in 1% BSA) for 2 hours, followed by 100 µl of HRP-conjugated streptavidin for 20 minutes. 100 µl of substrate solution (R&D Systems) was added to each well and the reaction was stopped by adding 50 µl of 2N $H_2SO_4$ (R&D Systems). The optical density of each well was determined immediately using a microplate reader (Molecular Devices) set to 450 nm.

**Multiplex human inflammatory cytokine assay**

Multiplex cytokine profiling was performed using the 12-Plex Human Inflammation Panel (Brightest Bio, HIF12). All reagents and assay procedures were performed according to the manufacturer's instructions. Specifically, after equilibration of all reagents to room temperature, serial dilutions of the cytokine standards and organoid medium samples (2-fold dilution) were prepared and added to designated wells. Plates were incubated for 3 hours at room temperature on an orbital shaker set to 650 rpm. Following incubation, wells were washed with supplied wash buffer, and 100 µl of biotinylated detection antibodies were added to each well. Plates were incubated for 1 hour under the same shaking conditions. The plate was then washed and incubated with 50 µl of Plasmonic Fluor (PF)-labeled streptavidin for 20 minutes at 650 rpm. Plates were then washed six times, and the fluorescence intensity of each well was determined using the Discovery Reader (Brightest Bio).

**Luciferase assay.**

Gaussia luciferase (Gluc) activity in conditioned medium from kidney organoids expressing SERCaMP, with the indicated treatments, was measured using the Gaussia Luciferase Glow Assay Kit (Thermo Fisher Scientific) following the manufacturer's instructions. Luminescence was detected with the GloMax-Multi Detection System (Promega). Raw light unit (RLU) values were normalized to protein concentration. Data were further normalized to control groups and expressed as fold changes.

**Organoid culture medium concentration**

For detecting UMOD protein in culture medium from kidney organoids, culture medium was collected and transferred to pre-chilled Amicon® Ultra centrifugal filter (50 kD) (Millipore) for protein enrichment. For each sample, 0.5 mL of medium was loaded into the filter unit and centrifuged at 4,000 × g at 4 °C until the retentate volume was reduced approximately 10-fold relative to the starting volume. The centrifugal filtration protocol was carried out according to the manufacturer's recommendations to maximize protein recovery and minimize membrane adsorption losses. The concentrated retentate was collected and mixed with SDS sample buffer for Western blot analysis.

**Statistics**

Statistical analyses were performed using GraphPad Prism (version 10) software. Data were presented as mean ± standard deviation (mean ± SD) or plots. For comparisons between two groups, a two-tailed Student's *t* test was applied. For comparisons among three or more groups, one-way analysis of variance (ANOVA) followed by post-hoc Tukey test was used to determine statistical significance. * $P < 0.05$, ** $P < 0.01$, *** $P < 0.001$, **** $P < 0.0001$. All experiments were independently repeated at least two to three times with consistent results.

**Study approval**

Human skin biopsies and paraffin-embedded kidney sections from ADTKD patients and their unaffected family members were obtained from the Wake Forest Cohort under a protocol approved by the Institutional Review Board of Wake Forest University School of Medicine. The written informed consent was obtained from all participants. The slides are stored and provided for the present study in a de-identified manner.

**Data availability**

All data needed to evaluate the conclusions in the paper are present in the paper or the Supplemental Materials. The generated raw fastq sequencing files and gene counts data in this study needed for reproducibility with publicly available tools cited in the Methods will be deposited in Gene Expression Omnibus [GEO] database, once the manuscript is accepted. Source data are provided with this paper.

**Acknowledgements**

We thank Dr. Sanjay Jain for an insightful discussion about our RNAseq data analysis. We thank the Genome Technology Access Center at the McDonnell Genome Institute at Washington University School of Medicine for help with genomic analysis. The Center is partially supported by NCI Cancer Center Support Grant #P30 CA91842 to the Siteman Cancer Center from the National Center for Research Resources (NCRR), a component of the National Institutes of Health (NIH), and NIH Roadmap for Medical Research. C.J.G is supported by American Society of Nephrology (ASN) Kidney Cure Ben J. Lipps Research Fellowship Program. Y.L.F is supported by the American Heart Association Postdoctoral Fellowship. F.U. is supported by NIH grants R01 DK132090 and P30 DK020579. M.L. is supported by Juvenile Diabetes Research Foundation 17-2013-410 and 2-SRA-2018-496-A-B, grants from the Finnish Diabetes Research Foundation and Grants 117044 and 333974 from the Academy of Finland. A.J.B. is supported by NIH grant R21DK106584 and Subaward Critical Path Institute FDA contract 75F40124C00106. S.S. is supported by National Science Foundation (CBET-2316285 and CBET-2331330) and the Congressionally Directed Medical Research Programs (HT94252310996). Y.M.C. is supported by NIH grants R01 2DK105056-6, R56DK138158A1, R21DK131557A1, R03DK106451 and K08DK089015, VA Merit I01BX006401A1, the Office of the Assistant Secretary of Defense for Health Affairs through the Peer Reviewed Medical Research Program under Award W81XWH-19-1-0320, George M. O'Brien Kidney Research Core Centers (NIH P30 DK114857; NIH P30 DK079337), Seed Grant from WashU Center of Regenerative Medicine, Mallinckrodt Challenge

Grant, Investigator Matching Micro Grant from WashU Center for Drug Discovery, and Faculty Scholar Award from the Children’s Discovery Institute of WashU and St. Louis Children's Hospital. Y.M.C. is a member of Washington University Institute of Clinical and Translational Sciences (UL1 TR000448) and Diabetes Research Center (NIH P30 DK020579).

### Table 1. List of Antibodies

| ANTIBODIES | SOURCE | HOST | CLONALITY | IDENTIFIER | DILUTION |
|---|---|---|---|---|---|
| **Primary antibodies** | | | | | |
| AMPKα | Cell Signaling Technology | Rabbit | Monoclonal | Cat #5831 | 1: 1000 |
| ATF6 | Novus Biologicals | Mouse | monoclonal | Cat #BP1-40256 | 1: 1000 |
| β-Actin-Peroxidase | Sigma-Aldrich | Mouse | monoclonal | Cat #A3854 | 1: 1000 |
| Cleaved Caspase-3 | Cell Signaling Technology | Rabbit | monoclonal | Cat #9664 | 1: 1000 |
| FLAG | Sigma-Aldrich | Mouse | monoclonal | Cat #F1804 | 1: 1000 |
| FOXO3a | Cell Signaling Technology | Rabbit | monoclonal | Cat #12829 | 1: 1000 |
| GRP78/BIP | Proteintech | Rabbit | polyclonal | Cat #11587-1-AP | 1: 1000 (WB)<br>1: 50 (IF) |
| HA | Sigma-Aldrich | Rabbit | polyclonal | Cat #H6908 | 1: 1000 |
| HA | Santa Cruz Biotechnology | Mouse | monoclonal | Cat #sc-7392 | 1: 1000 |
| IP3R1 | Cell Signaling Technology | Rabbit | monoclonal | Cat #8568 | 1: 1000 (WB)<br>1: 50 (IF) |
| LC3B | Cell Signaling Technology | Rabbit | polyclonal | Cat #2775 | 1: 1000 |
| MANF | Abnova | Rabbit | polyclonal | Cat #PAB13301 | 1: 1000 (WB)<br>1: 50 (IF) |
| MANF | R&D Systems | Goat | polyclonal | Cat #AF3748 | 1: 1000 |
| Phospho-AMPKα (T172/T183) | Cell Signaling Technology | Rabbit | monoclonal | Cat #50081 | 1: 1000 |
| SQSTM1/p62 | Cell Signaling Technology | Rabbit | polyclonal | Cat #5114 | 1: 1000 |
| TRIB3 | Cell Signaling Technology | Rabbit | monoclonal | Cat #43043 | 1: 1000 |
| TRIB3 | Proteintech | Rabbit | polyclonal | Cat #13300-1-AP | 1: 1000 (WB)<br>1: 50 (IF) |
| UMOD | Alfa Aesar | Rabbit | polyclonal | Cat #J65429 | 1: 1000 |
| UMOD | RayBiotech | Mouse | monoclonal | Cat # 119-13298 | 1: 50 (IF) |
| UMOD | R&D Systems | Sheep | polyclonal | Cat #AF5144 | 1: 50 (IF) |
| UMOD Biotinylated | R&D Systems | Sheep | polyclonal | Cat #BAF5144 | 4 µg/sample (TAL isolation) |
| **Secondary antibodies** | | | | | |
| Anti-Rabbit IgG, secondary antibody, Alexa Fluor™ 488 | Invitrogen | Donkey | polyclonal | Cat #A-21206 | 1:500 |
| Anti-Sheep IgG, secondary antibody, Alexa Fluor™ 594 | Invitrogen | Donkey | polyclonal | Cat #A-11016 | 1:500 |
| Secondary anti-Rabbit IgG, HRP-linked | Cell Signaling Technology | Goat | polyclonal | Cat #7074S | 1:5000 |
| Secondary anti-Mouse IgG, HRP-linked | Cell Signaling Technology | Goat | polyclonal | Cat #7076S | 1:5000 |

### Table 2. List of Reagents and Assays

| | SOURCE | IDENTIFIER |
|---|---|---|
| **Reagents** | | |
| 4',6-diamidino-2-phenylindole (DAPI) | Invitrogen | Cat #D1306 |
| Accutase cell dissociation reagent | Gibco | Cat #A1110501 |
| APEL2 medium | Stem Cell Technologies | Cat #5270 |
| Blasticidin S HCl | Gold Biotechnology | Cat #B-800-25 |
| CHIR 99021 | Tocris Bioscience | Cat #4423 |
| Clarity Max Western ECL substrate | Bio-Rad Laboratories | Cat # 1705062 |
| DC Protein Assay | Bio-Rad Laboratories | Cat #5000111 |
| DMEM, high glucose | Gibco | Cat #11965092 |
| DMEM/F-12 | Gibco | Cat #11320082 |
| Doxycycline hyclate | Sigma-Aldrich | Cat #33429 |
| DPBS | Gibco | Cat #14190250 |
| Dynabeads Biotin Binder | Invitrogen | Cat #11047 |
| Fast SYBR Green Master Mix | Applied Biosystems | Cat #4385612 |
| Fetal Bovine Serum | Gibco | Cat #A5256801 |
| Fluoromount-G Mounting Medium | Invitrogen | Cat #00-4958-02 |
| Heparin sodium | Sigma-Aldrich | Cat #H3150 |
| Lipofectamine 3000 Transfection Reagent | Invitrogen | Cat #L3000015 |
| Matrix | Corning | Cat #354277 |
| mTeSR Plus medium | Stem Cell Technologies | Cat #1000276 |
| Protease and Phosphatase Inhibitor Cocktail | Sigma-Aldrich | Cat # PPC2020 |
| Protein A/G PLUS-Agarose | Santa Cruz Biotechnology | Cat #sc-2003 |
| Puromycin dihydrochloride | Sigma-Aldrich | Cat #P9620 |
| Recombinant human FGF9 protein | R&D Systems | 273-F9-025 |
| RIPA buffer | Cell Signaling Technology | Cat #9806 |
| Stop Solution: 2 N H2SO4 | R&D Systems | Cat #DY994 |
| Substrate Solution | R&D Systems | Cat #DY999 |
| Tissue-Tek OCT Compound | Sakura Finetek | Cat #4583 |
| Xestospongin C | Sigma-Aldrich | Cat #X2628 |
| Y-27632 Dihydrochloride | Tocris Bioscience | Cat #1254 |
| **Assays** | | |
| Gaussia Luciferase Glow Assay Kit | Thermo Fisher Scientific | Cat #16160 |
| Gibson Assembly Master Mix | NEB | Cat #E2611S |
| Human IL-6 DuoSet ELISA kit | R&D Systems | Cat #DY206 |
| KOD-Plus-Mutagenesis Kit | TOYOBO | Cat #SMK-101 |
| NucleoBond Xtra Midi kit for endotoxin-free plasmid DNA | Macherey-Nagel | Cat #740410.50 |
| PrimeScript RT Reagent Kit | Takara Bio | Cat #RR047A |
| RNeasy UCP Micro Kit | QIAGEN | Cat #73934 |

### Table 3. List of Primers for Kidney Organoids

| Primers | SOURCE | Sequence (5′-3′) |
|---|---|---|
| Genotyping primers for *UMOD* | Integrated DNA Technologies | Forward: CATGTGTCAATGTGGTGGGC<br>Reverse: TAGCCCTCCCCGTACTCG |
| qPCR primers for *ATG5* | Integrated DNA Technologies | Forward: AAAGATGTGCTTCGAGATGTGT<br>Reverse: CACTTTGTCAGTTACCAACGTCA |
| qPCR primers for *ATG7* | Integrated DNA Technologies | Forward: CTGCCAGCTCGCTTAACATTG<br>Reverse: CTTGTTGAGGAGTACAGGGTTTT |
| qPCR primers for *BECN1* | Integrated DNA Technologies | Forward: CCATGCAGGTGAGCTTCGT<br>Reverse: GAATCTGCGAGAGACACCATC |
| qPCR primers for *CCL2* | Integrated DNA Technologies | Forward: CAGCCAGATGCAATCAATGCC<br>Reverse: TGGAATCCTGAACCCACTTCT |
| qPCR primers for *GABARAPL1* | Integrated DNA Technologies | Forward: ATGAAGTTCCAGTACAAGGAGGA<br>Reverse: GCTTTTGGAGCCTTCTCTACAAT |
| qPCR primers for *GAPDH* | Integrated DNA Technologies | Forward: GTCTCCTCTGACTTCAACAGCG<br>Reverse: ACCACCCTGTTGCTGTAGCCAA |
| qPCR primers for *ICAM1* | Integrated DNA Technologies | Forward: ATGCCCAGACATCTGTGTCC<br>Reverse: GGGGTCTCTATGCCCAACAA |
| qPCR primers for *IL1B* | Integrated DNA Technologies | Forward: AGCTACGAATCTCCGACCAC<br>Reverse: CGTTATCCCATGTGTCGAAGAA |
| qPCR primers for *IL6* | Integrated DNA Technologies | Forward: ACTCACCTCTTCAGAACGAATTG<br>Reverse: CCATCTTTGGAAGGTTCAGGTTG |
| qPCR primers for *ITPR1/IP3R1* | Integrated DNA Technologies | Forward: GCGGAGGGATCGACAAATGG<br>Reverse: TGGGACATAGCTTAAAGAGGCA |
| qPCR primers for *ITPR2/IP3R2* | Integrated DNA Technologies | Forward: CACCTTGGGGTTAGTGGATGA<br>Reverse: CTCGGTGTGGTTCCCTTGT |
| qPCR primers for *ITPR3/IP3R3* | Integrated DNA Technologies | Forward: CCAAGCAGACTAAGCAGGACA<br>Reverse: ACACTGCCATACTTCACGACA |
| qPCR primers for *MAP1LC3B* | Integrated DNA Technologies | Forward: GATGTCCGACTTATTCGAGAGC<br>Reverse: TTGAGCTGTAAGCGCCTTCTA |
| qPCR primers for *RYR1* | Integrated DNA Technologies | Forward: CATCAGCACGACATGAGCTT<br>Reverse: CCCACACCATGTAGCAGTTG |
| qPCR primers for *RYR2* | Integrated DNA Technologies | Forward: TGCAAGACTCACCGAAGATG<br>Reverse: CCACCCAGACATTAGCAGGT |
| qPCR primers for *RYR3* | Integrated DNA Technologies | Forward: CTGACGTCCATCTTTGAGCA<br>Reverse: CAAGGGAGTAGAGGCTGCAC |
| qPCR primers for *TNFA* | Integrated DNA Technologies | Forward: CCTCTCTCTAATCAGCCCTCTG<br>Reverse: GAGGACCTGGGAGTAGATGAG |
| qPCR primers for *ULK1* | Integrated DNA Technologies | Forward: AGCACGATTTGGAGGTCGC<br>Reverse: GCCACGATGTTTTCATGTTTCA |

### Table 4. List of Plasmids

| Plasmid | SOURCE | IDENTIFIER |
|---|---|---|
| pcDNA3.1-FLAG | This paper | N/A |
| pcDNA3.1-FLAG-AMPKα1 | This paper | N/A |
| pcDNA3.1-FLAG-AMPKα1-AL deletion | This paper | N/A |
| pcDNA3.1-FLAG-AMPKα1-C terminus deletion | This paper | N/A |
| pcDNA3.1-FLAG-AMPKα1-N terminus deletion | This paper | N/A |
| pcDNA3.1-FLAG-AMPKα2 | This paper | N/A |
| pcDNA3.1-FLAG-AMPKα2-AL deletion | This paper | N/A |
| pcDNA3.1-FLAG-AMPKα2-C terminus deletion | This paper | N/A |
| pcDNA3.1-FLAG-AMPKα2-N terminus deletion | This paper | N/A |
| pcDNA3.1-HA | This paper | N/A |
| pcDNA3.1-HA-TRIB3 | This paper | N/A |
| pLenti6.3-PGK-SERCaMP | This paper | N/A |
| pLKO.1 | Addgene | Cat #10878 |
| pLVX-CMV-MANF | This paper | N/A |
| pLVX-TRE3G-MANF-PGK-rtTA-P2A-Blast | This paper | N/A |
| pMD2.G | Addgene | Cat #12259 |
| psPAX2 | Addgene | Cat #12260 |

### Supplemental Figures

A

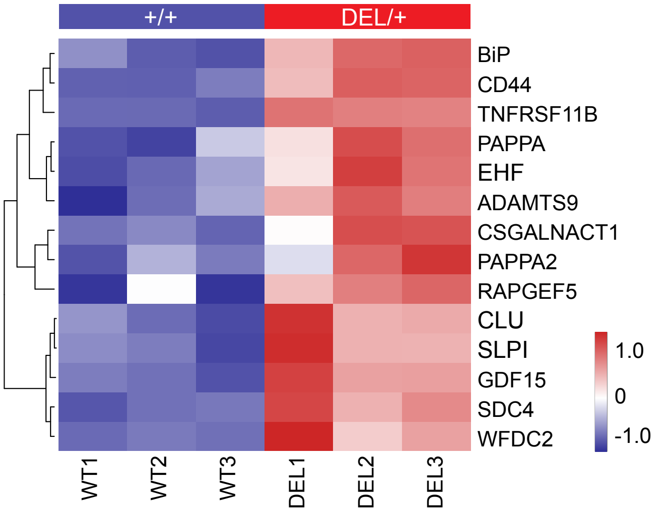


B

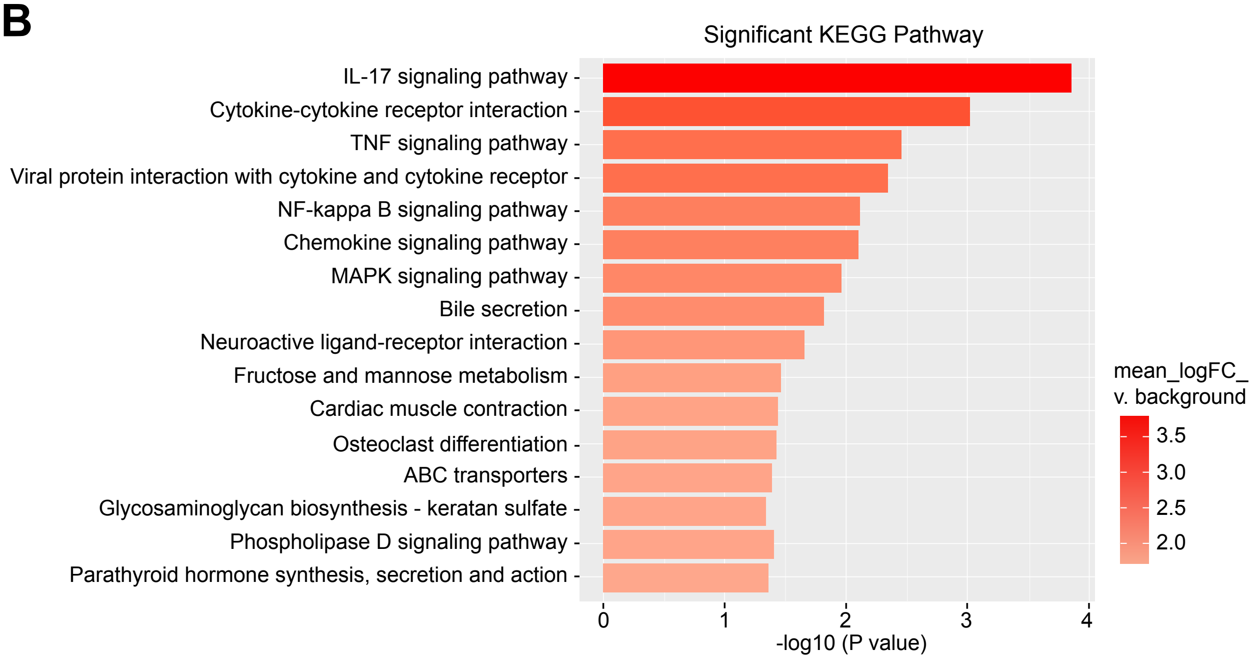


**Supplemental Figure 1. RNA-seq analysis of *UMOD* $^{DEL/+}$ vs. WT TALs isolated from kidney organoids.** (**A**) Heatmap of upregulated TAL injury state markers, as identified in HKAv2 atlas, in the mutant TALs compared with WT TALs. Benjamini-Hochberg adjusted $P \leq 0.05$. GAGE analysis. (**B**) KEGG pathway analysis of the log2-fold changes in the isolated mutant TALs compared with WT TALs. GAGE analysis.

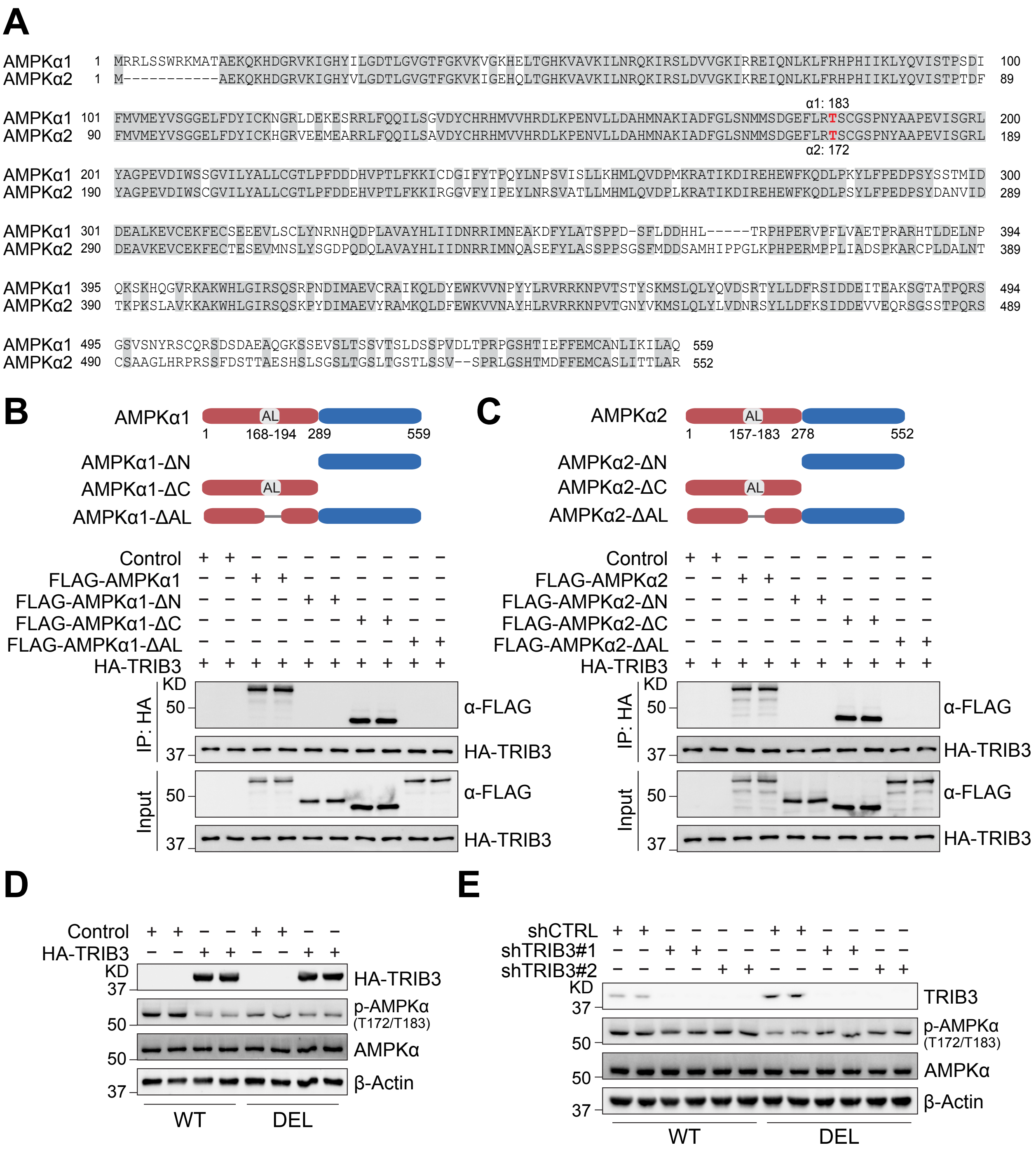
A
AMPKα1
AMPKα2
α1: 183
α2: 172
B
AMPKα1
AL
1
168-194
289
559
AMPKα1-ΔN
AMPKα1-ΔC
AMPKα1-ΔAL
Control
FLAG-AMPKα1
FLAG-AMPKα1-ΔN
FLAG-AMPKα1-ΔC
FLAG-AMPKα1-ΔAL
HA-TRIB3
KD
IP: HA
Input
α-FLAG
HA-TRIB3
C
AMPKα2
157-183
278
552
AMPKα2-ΔN
AMPKα2-ΔC
AMPKα2-ΔAL
FLAG-AMPKα2
FLAG-AMPKα2-ΔN
FLAG-AMPKα2-ΔC
FLAG-AMPKα2-ΔAL
D
p-AMPKα
(T172/T183)
AMPKα
β-Actin
WT
DEL
E
shCTRL
shTRIB3#1
shTRIB3#2
TRIB3

**Supplemental Figure 2. TRIB3 binds to activation loop (AL) domain of AMPKα.** (**A**) Amino acid sequence alignment of human AMPKα1 and AMPKα2, with conserved residues highlighted. T183 in AMPKα1 or T172 in AMPKα2 within the AL domain is required for AMPK activation. (**B**) *Upper panel*, schematic representation of AMPKα1 domain. *Lower panel*, HEK 293T cells were co-transfected with expression plasmids of control, FLAG-full length AMPKα1 or deletion mutants, together with HA-TRIB3. Cell extracts were IP with anti-HA Ab and blotted with anti-FLAG Ab. (**C**) *Upper panel*, schematic representation of AMPKα2 domain. *Lower panel*, HEK 293T cells were co-transfected with expression plasmids of control, FLAG-full length AMPKα2 or deletion mutants, with HA-TRIB3. Cell extracts were IP with anti-HA Ab and blotted with anti-FLAG Ab. AMPKα truncated mutants include full-length protein lacking the N-terminal domain (ΔN), the C-terminal domain (ΔC) and activation loop (ΔAL). (**D**) Western blot analysis of TRIB3, p-AMPKα (T172/T183) and AMPKα in cell lysates of WT and DEL cells transfected with either Control or HA-TRIB3 plasmids for 48 hours. (**E**) Western blot analysis of TRIB3, p-AMPKα (T172/T183) and AMPKα in cell lysates of WT and DEL cells transduced with Control or shTRIB3 for 72 h**.**

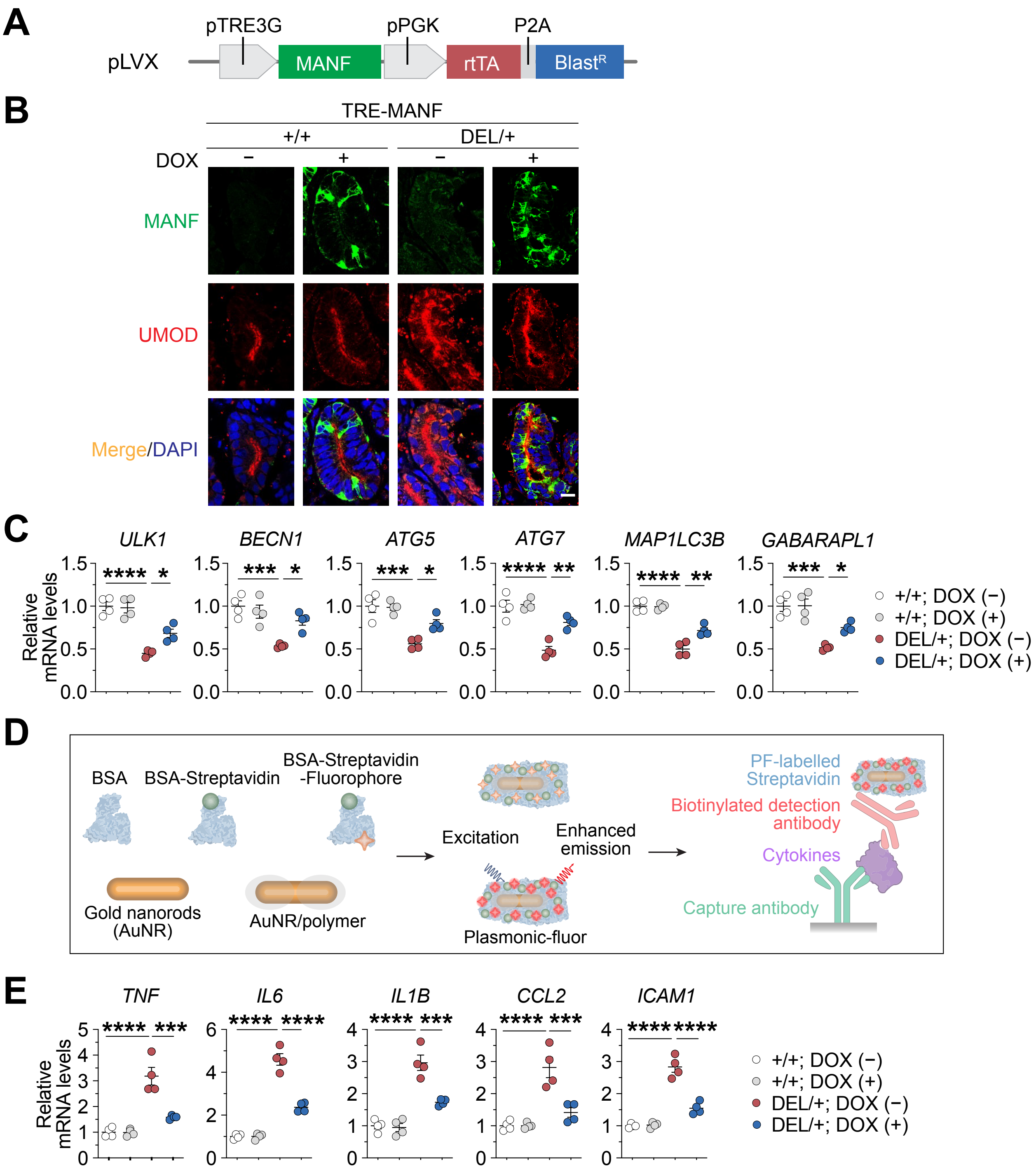
A
pLVX
pTRE3G
MANF
pPGK
rtTA
P2A
BlastR
B
TRE-MANF
+/+
DEL/+
DOX
−
+
MANF
UMOD
Merge/DAPI
C
ULK1
BECN1
ATG5
ATG7
MAP1LC3B
GABARAPL1
Relative mRNA levels
+/+; DOX (−)
+/+; DOX (+)
DEL/+; DOX (−)
DEL/+; DOX (+)
D
BSA
BSA-Streptavidin
BSA-Streptavidin -Fluorophore
Gold nanorods (AuNR)
AuNR/polymer
Excitation
Enhanced emission
Plasmonic-fluor
PF-labelled Streptavidin
Biotinylated detection antibody
Cytokines
Capture antibody
E
TNF
IL6
IL1B
CCL2
ICAM1
Relative mRNA levels
+/+; DOX (−)
+/+; DOX (+)
DEL/+; DOX (−)
DEL/+; DOX (+)

**Supplemental Figure 3. Lentiviral vector-mediated inducible MANF overexpression after the kidney organoid differentiation stimulates autophagic degradation of mutant UMOD.** (**A**) Diagram of Tet-inducible MANF expression system. TRE3G: tetracycline responsive element 3G. PGK promoter: phosphoglycerate kinase 1 promoter. rtTA: reverse tetracycline-controlled transactivator. P2A: porcine teschovirus-1 2A peptide. With DOX, rtTA binds to TRE3G and initiates transcription of the human MANF cDNA. (**B**) Representative co-IF images of WT and mutant kidney organoids without or with the administration of DOX from days 22-30, stained for MANF (green) and UMOD (red) with nuclei counterstain (blue). Scale bar, 10 μm. (**C**) Transcript analysis of a panel of autophagy-related genes in WT and mutant organoid-derived TALs treated without DOX or with DOX from days 22-30. (**D**) Schematic illustration of plasmon-enhanced fluorescence-linked immunosorbent multiplex assay (p-FLISA). The structure of plasmonic-fluor nanosensor comprises a plasmonically active core (an AuNR), a polymer shell that acts as a spacer layer, light emitters, and a universal biorecognition element (biotin). Bovine serum albumin (BSA) serves as a crucial design element to assemble all components into the functional nanoconstruct and to minimize non-specific binding. Plasmonic-fluor, as an "add-on" biolabel, enhances fluorescence intensity and signal-to-noise ratio dramatically in p-FLISA without altering the existing workflow. (**E**) Relative mRNA levels of pro-inflammatory cytokines in WT and mutant organoid-isolated TALs administered without or with DOX from days 22-30. * $P < 0.05$; ** $P < 0.01$; *** $P < 0.001$; **** $P < 0.0001$.

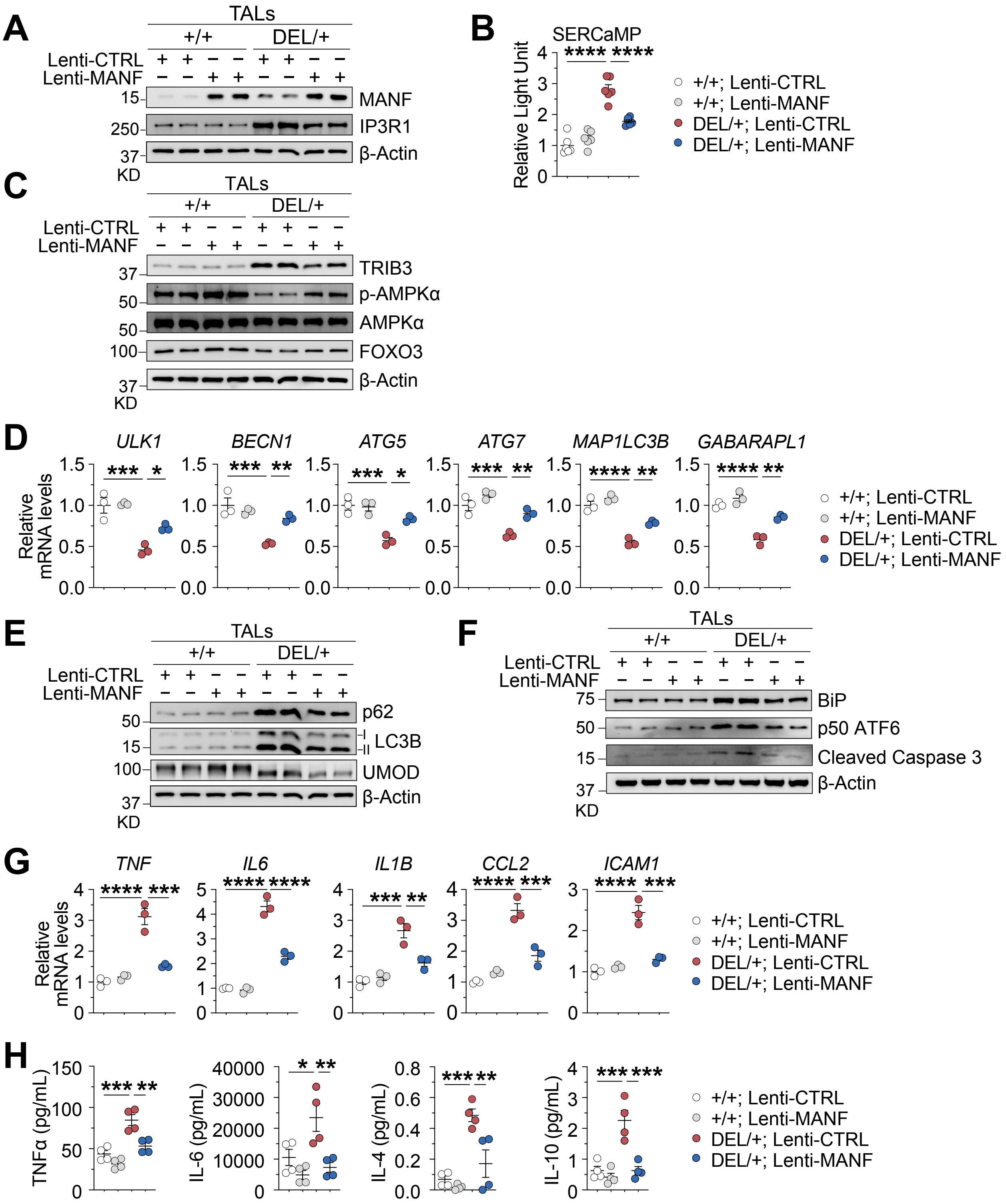

A
TALs
+/+
DEL/+
Lenti-CTRL
Lenti-MANF
MANF
IP3R1
β-Actin
KD
B
SERCaMP
Relative Light Unit
+/+; Lenti-CTRL
+/+; Lenti-MANF
DEL/+; Lenti-CTRL
DEL/+; Lenti-MANF
C
TRIB3
p-AMPKα
AMPKα
FOXO3
D
ULK1
BECN1
ATG5
ATG7
MAP1LC3B
GABARAPL1
Relative mRNA levels
E
p62
LC3B
UMOD
F
BiP
p50 ATF6
Cleaved Caspase 3
G
TNF
IL6
IL1B
CCL2
ICAM1
H
TNFα (pg/mL)
IL-6 (pg/mL)
IL-4 (pg/mL)
IL-10 (pg/mL)

**Supplemental Figure 4. Lentivirus expressing MANF decreases mutant UMOD accumulation and mitigates ER stress-associated inflammation in *UMOD* $^{DEL/+}$ TALs.** WT and DEL/+ organoids were infected with control or lentivirus overexpressing MANF from days 22-30. At the end of the virus infection, TALs were isolated from the WT or mutant organoids, and medium were collected. (**A, C, E and F**) IB analysis of indicated proteins in TALs treated without or with MANF. (**B**) Luciferase activity in the culture medium of WT or mutant kidney organoids, which were differentiated from stable SERCaMP-expressing WT or mutant hiPSCs respectively, without or with MANF treatment. (**D**) Q-PCR analysis of a panel of autophagy-related genes in WT or mutant TALs in the absence or presence of MANF treatment. (**G**) Relative mRNA levels of pro-inflammatory cytokines in WT or DEL/+ TALs in the absence or presence of MANF treatment. (**H**) p-FLISA of pro-inflammatory cytokine levels in the medium of WT or mutant kidney organoids, without or with MANF treatment. Data are shown as mean ± SD. * $P < 0.05$; ** $P < 0.01$; *** $P < 0.001$; **** $P < 0.0001$.
